# PKMYT1 inhibition induces DNA damage and synergizes with immune checkpoint blockade in *CCNE1*-amplified gastroesophageal adenocarcinoma

**DOI:** 10.64898/2025.12.17.694745

**Authors:** Sungjoo Jang, Lawrence W. Wu, Jimyung Park, Sho Hangai, Cameron Shapiro, C. Gary Marshall, Luke L. Cai, Benjamin Izar, Adam J. Bass, Ryan H. Moy

## Abstract

**Purpose:** *CCNE1* amplification, found in approximately 10% of gastroesophageal adenocarcinoma (GEA), drives chromosomal instability and is associated with an immune-cold tumor microenvironment. Recent studies suggest that PKMYT1 inhibition is synthetic lethal in *CCNE1*-amplified cancers; however, its role in GEA is largely uncharacterized. We investigated the therapeutic activity of PKMYT1 inhibition in *CCNE1*-amplified GEA and its potential combination with immunotherapy.

**Experimental Design:** We evaluated lunresertib (RP-6306), a selective first-in-class PKMYT1 inhibitor, in *CCNE1*-amplified and wild-type GEA cell lines to define its effects on cell viability, DNA damage, and gene expression changes. To investigate the role of PKMYT1 blockade on immune microenvironment modulation, we used a syngeneic murine model of CCNE1;*Trp53*^-/-^ GEA xenografts treated with lunresertib, anti-PD-1, or the combination.

**Results:** Lunresertib was selectively cytotoxic in *CCNE1*-amplified GEA cell lines in cell viability and clonogenic assays. Mechanistically, treatment with lunresertib induced DNA damage, evidence by increased γH2AX expression and DNA micronuclei formation, and activated pro-inflammatory signaling pathways by gene set enrichment analysis. In a *CCNE1*-amplified GEA murine model, PKMYT1 inhibition promoted T cell infiltration, reduced myeloid cells, and synergized with anti-PD-1 therapy to induce tumor regression.

**Conclusions:** Our results suggest that *CCNE1* amplification is a potentially actionable target in GEA and support the development of combination therapeutic strategies utilizing PKMYT1 inhibition and PD-1 blockade.

**Translational Relevance:** Gastroesophageal adenocarcinoma (GEA) is an aggressive malignancy with poor prognosis and few actionable molecular alterations. *CCNE1* amplification is present in approximately 10% of patients with GEA and represents a potential therapeutic vulnerability through synthetic lethality with PKMYT1 inhibition. We demonstrate that targeting *CCNE1*-amplified GEA through PKMYT1 inhibition with lunresertib is selectively cytotoxic, induces DNA damage, and activates inflammatory signaling pathways. In a syngeneic murine model of *CCNE1*-amplified gastric cancer, we further identified that PKMYT1 inhibition induced T cell infiltration and tumor microenvironment remodeling, and synergized with anti-PD-1 therapy to drive tumor regression. Together, these results support further development of PKMYT1 inhibition in combination with PD-1 blockade for *CCNE1*-amplified GEA.

## Introduction

Gastroesophageal adenocarcinoma (GEA)—including gastric cancer (GC), gastroesophageal junction (GEJ) adenocarcinoma, and esophageal adenocarcinoma (EA)—is a major global health burden with nearly one million new diagnoses annually, ranking as the fourth leading cause of cancer-related mortality (1). Large-scale genomic sequencing such as through The Cancer Genome Atlas (TCGA) have defined recurrent genomic alterations in GEA and identified four main molecular subtypes: chromosomal instability (CIN), genomically stable (GS), microsatellite instability (MSI), and Epstein-Barr virus-positive (2,3). Despite this molecular framework, the number of validated targetable alterations and predictive biomarkers in GEA remains limited, apart from microsatellite instability, programmed cell death ligand 1 (PD-L1), human epidermal growth factor receptor 2 (HER2), and more recently Claudin 18.2 (CLDN18.2) (4,5). Immunotherapy with anti-PD-1 inhibitors such as nivolumab, pembrolizumab, and tislelizumab are now approved in combination with first-line platinum-based chemotherapy for metastatic disease (6–8). Even with the advances of modern chemotherapy and immunotherapy, the long-term prognosis for advanced GEA remains poor with a five-year survival rate of less than 10% (9). Therefore, there is a critical need to identify additional clinically relevant therapeutic targets and enhance immunotherapy responsiveness.

One genomic alteration that may modulate the tumor-immune microenvironment and represent a therapeutic vulnerability is amplification of the cell cycle regulator Cyclin E (*CCNE1*). *CCNE1* is amplified in multiple cancers including approximately 10% of esophageal and gastric cancers (10–12). Functionally, *CCNE1* promotes cell cycle progression through the G1/S phase as it binds and activates CDK2, initiating DNA replication. Aberrant *CCNE1* amplification leads to premature CDK2 activation, resulting in replication stress, aneuploidy, and chromosome instability (CIN), which are key drivers of oncogenesis and tumor progression (13). While it is still unclear whether *CCNE1* amplification is directly causative of CIN, the high levels of replication stress due to premature mitotic entry produces chromosome segregation defects and genomic instability, raising CIN levels. (14) In patients with gastric cancer, *CCNE1* amplification and CIN have been associated with increased frequency of liver metastasis and poor overall survival (10,15). *CCNE1* is also frequently co-amplified with *ERBB2* and promotes resistance to HER2-targeted therapy (16). While CIN can activate innate immune sensing pathways, it can also promote immune evasion through several different mechanisms (17–19). Indeed, analysis of TCGA data demonstrated that CIN gastric cancers with *CCNE1* amplification are enriched for an immune-cold phenotype with reduced CD8+ T cell infiltration (20). Hence, targeting *CCNE1* amplification and/or CIN in GEA could represent a novel strategy to reprogram the tumor-immune microenvironment and augment immunotherapy responsiveness.

Although *CCNE1* amplification is not currently directly druggable, *CCNE1* amplification has been regarded as a potential target for synthetic lethality approaches. Recent synthetic lethality screens identified that *CCNE1*-amplified tumors are vulnerable to loss of membrane-associated tyrosine- and threonine-specific cdc2-inhibitory kinase (PKMYT1), a member of the Wee1 G2 checkpoint kinase family (21). PKMYT1 negatively regulates CDK1 through inhibitory phosphorylation on Thr14 and its sequestration in the cytoplasm (22–24). In CCNE1 overexpressing cells, PKMYT1 inhibition leads to CDK1 hyperactivation, unscheduled mitotic entry, catastrophic DNA damage, and cell death (21). Lunresertib (RP-6306) is a selective, first-in-class, oral PKMYT1 inhibitor which is currently in early-phase clinical trials and has shown clinical activity as monotherapy or in combination with targeted therapy (such as ATR inhibitors) or chemotherapy (25). However, the effect of PKMYT1 inhibition in *CCNE1*-amplified GEA has not been explored. Additionally, in recent years, immunotherapy has emerged as a new standard of care for several malignancies including GEA and has demonstrated increased clinical benefit in some populations (26). As DNA damage and extranuclear DNA may activate innate immune signaling, we hypothesized that PKMYT1 inhibition may extend beyond tumor-intrinsic proliferation control by modulating the tumor immune microenvironment and potentiating immunotherapy responses in CCNE1-overexpressing tumors. In this study, we investigated the efficacy of PKMYT1 inhibition and combinations with immune checkpoint blockade in preclinical models of *CCNE1-*amplified GEA.

## Results

### PKMYT1 inhibition is selectively cytotoxic in *CCNE1*-amplified GEA

Lunresertib has been shown to be synthetically lethal in *CCNE1*-amplified cancer cells such as ovarian, endometrial, and breast cancers (27,28), but its antitumor activity in GEA has not been well defined. To determine whether *CCNE1* amplification sensitizes GEA to PKMYT1 inhibition, we utilized two-dimensional cell lines derived from three-dimensional GEA patient-derived organoids (PDOs). We validated that these PDO-derived cell lines (MW5NT and MW5FJ) showed increased *CCNE1* expression compared to a *CCNE1* wild-type (WT) GC cell line (SNU668) by immunoblot (Figure 1A) and qRT-PCR (Supplementary Figure 1A). *CCNE1* amplification is an important driver of aneuploidy and CIN, which is characterized by chromosome mis-segregation errors that can lead to packaging of excess chromatin into DNA micronuclei (29). We measured CIN in *CCNE1* WT and amplified cell lines by quantifying DNA micronuclei. As expected, *CCNE1-*amplified MW5NT cells demonstrated a significantly higher percentage of cells bearing micronuclei compared to SNU668, consistent with a CIN phenotype in MW5NT cells (Figure 1B). After verifying elevated levels of *CCNE1*, we evaluated lunresertib sensitivity in MW5NT and SNU668 cells using cell proliferation and clonogenic assays. MW5NT cells demonstrated increased sensitivity to lunresertib with a IC50 of approximately 250nM, while SNU668 showed a IC50 of 5µM (Figure 1C). Clonogenic assays corroborated these findings, with *CCNE1*-amplified MW5NT showing reduced colony formation ability compared to SNU668 cells at equivalent lunresertib concentrations (Figure 1D). Similarly, lunresertib treatment suppressed colony formation rate in *CCNE1*-amplified MW5FJ cells (Supplementary Figure 1B). Taken together, these data demonstrate that PKMYT1 inhibition reduces proliferation and induces cytotoxicity selectively in *CCNE1*-amplified GEA.

**Figure 1.**
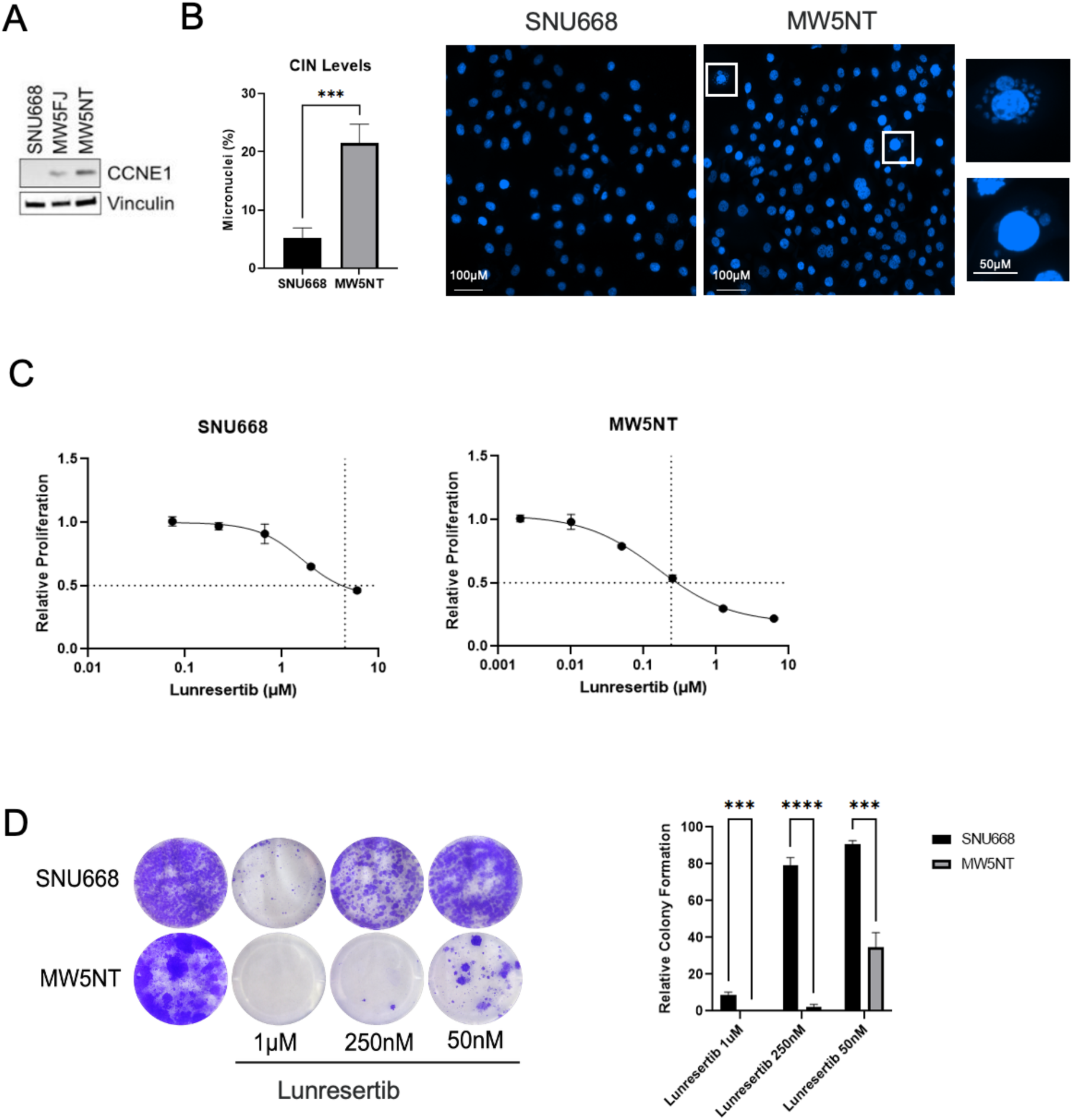
PKMYT1 inhibition is selectively cytotoxic in *CCNE1-*amplified GEA. **A.** CCNE1 expression in *CCNE1* WT (SNU668) and *CCNE1*-amplified (MW5FJ, MW5NT) by immunoblot. **B.** CIN levels of SNU668 and MW5NT cells quantified by counting micronuclei-bearing cells with representative images of micronuclei. **C.** Relative proliferation of SNU668 and MW5NT cells treated with indicated concentrations of lunresertib. **D.** Clonogenic assay of SNU668 and MW5NT cells treated with indicated concentrations of lunresertib. ***p < 0.001; ****p < 0.0001, Student’s t-test.

### PKMYT1 inhibition induces DNA damage in *CCNE1*-amplified GEA

During the G2 phase of the cell cycle, PKMYT1 phosphorylates Thr14 to inactivate the Cyclin B1-CDK1 complex to prevent premature mitosis, particularly in response to replication stress. In *CCNE1*-amplified cells, where replication stress is high, inhibition of PKMYT1 can induce forced mitotic entry and further errors in mitosis, resulting in mitotic catastrophe and cell death. To test if PKMYT1 inhibition leads to DNA damage in *CCNE1*-amplifed GEA models, we analyzed levels of γH2AX, a molecular marker of DNA double-strand breaks, following treatment with 1 µM of lunresertib for 24 hours (H) and 48H. While *CCNE1*-amplified MW5NT cells showed high levels of γH2AX protein expression after lunresertib treatment, there was minimal induction of γH2AX expression in *CCNE1* WT SNU668 cells (Figure 2A). Validating these findings, lunresertib treatment resulted in increased nuclear γH2AX staining by immunofluorescence in MW5NT cells relative to SNU668 (Figure 2B-C). Finally, we observed a higher proportion of MW5NT cells bearing DNA micronuclei after lunresertib treatment (Figure 2D). These results show that PKMYT1 inhibition selectively induces DNA damage and exacerbates CIN in *CCNE1*-amplified GEA.

**Figure 2.**
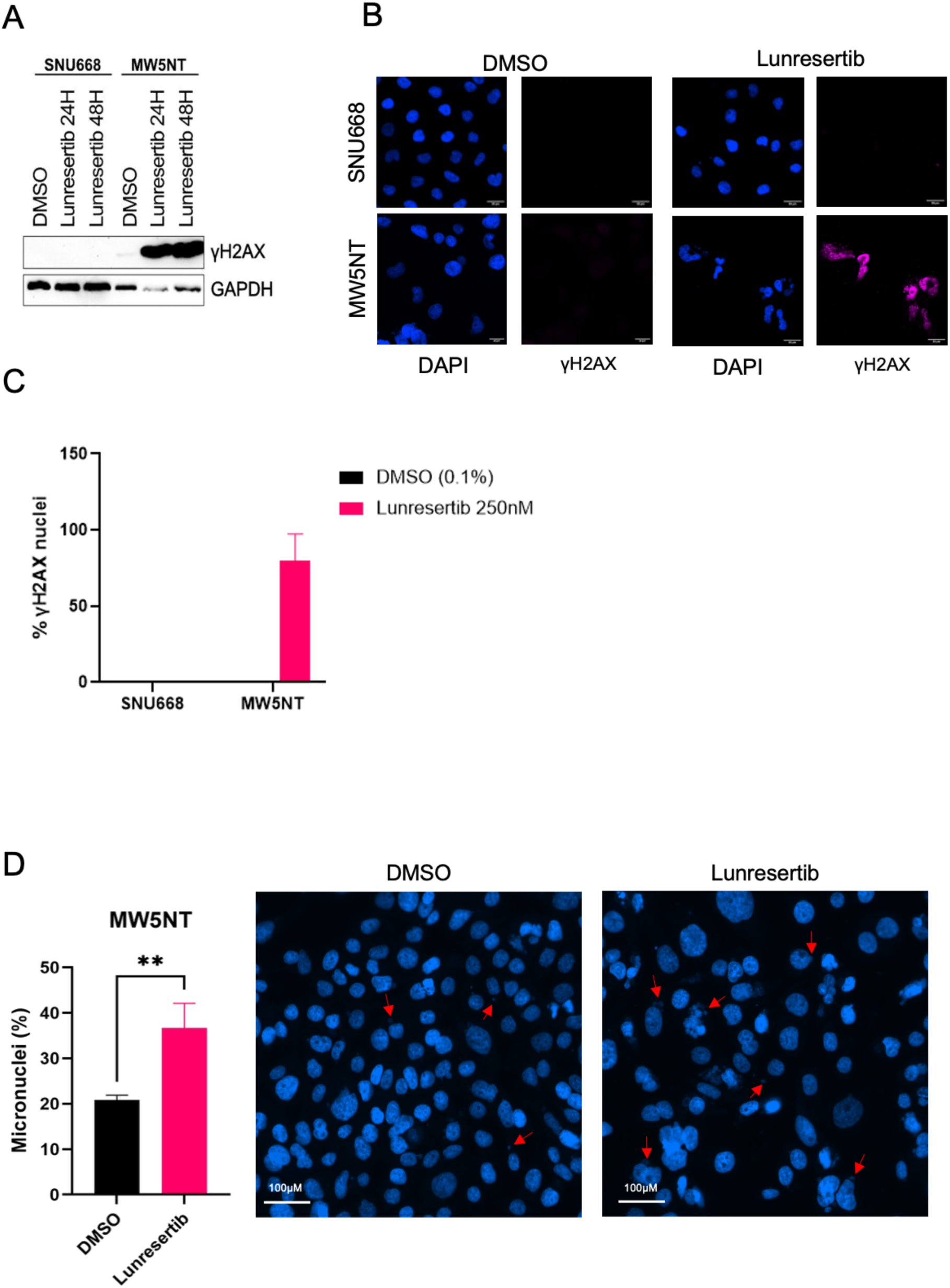
PKMYT1 inhibition induces DNA damage in *CCNE1-*amplified GEA. **A.** γH2AX levels of SNU668 and MW5NT cells treated with lunresertib or vehicle control by immunoblot. **B.** Nuclear γH2AX expression inSNU668 and MW5NT cells through immunofluorescence. **C.** Quantification of nuclear γH2AX expression in SNU668 and MW5NT cells treated lunresertib or vehicle control. **D.** CIN levels of MW5NT treated with DMSO vehicle control or lunresertib quantified by micronuclei-bearing cells. Representative images are shown. **p <0.01; Student’s t test.

### PKMYT1 inhibition induces pro-inflammatory signaling in *CCNE1*-amplified GEA

Next, we evaluated the effect of PKMYT1 inhibition and the DNA damage response on the transcriptional profile of *CCNE1*-amplified GEA. We exposed MW5NT cells to 1 µM of lunresertib for 48H and performed RNA sequencing (RNA-seq) to quantify gene expression changes (Figure 3A). To define biological processes associated with these transcriptional changes, we performed gene-set enrichment analysis on differentially expressed genes using the MSigDB Hallmark gene sets. Notably, the top profiles enriched in lunresertib-exposed cells included hallmark inflammatory and immune pathways including Tissue Necrosis Factor (TNF)-alpha signaling via NFκB, inflammatory response, interferon alpha response, interferon gamma response, and IL-6-JAK-STAT3 signaling (Figure 3B, C). Additionally, cell cycle and cell death programs were enriched, including upregulation of apoptosis and downregulation of E2F targets and G2M checkpoint (Figure 3B, C). We also examined KEGG pathways, with the top enriched pathway being cytokine-cytokine receptor interactions (Supplementary Figure 2A). From RNA sequencing data, select inflammatory genes such as interferon-stimulated gene 15 (*ISG15*), and C-X-C motif chemokine ligand 10 (*CXCL10*) were upregulated in lunresertib-treated MW5NT cells (Supplementary Figure 2B). We validated a subset of induced genes by quantitative reverse transcription PCR (qRT-PCR), confirming higher expression of *IFNβ1* and *ISG15* after lunresertib treatment in MW5NT cells compared to vehicle-treated controls (Figure 3D). Moreover, *CXCL5* and *CXCL10* expression was significantly increased after lunresertib treatment in MW5NT cells (Figure 3E). Thus, transcriptional profiling suggests that in addition to cell cycle dysregulation and cell death, PKMYT1 inhibition induces robust inflammatory response in *CCNE1*-amplified GEA.

**Figure 3.**
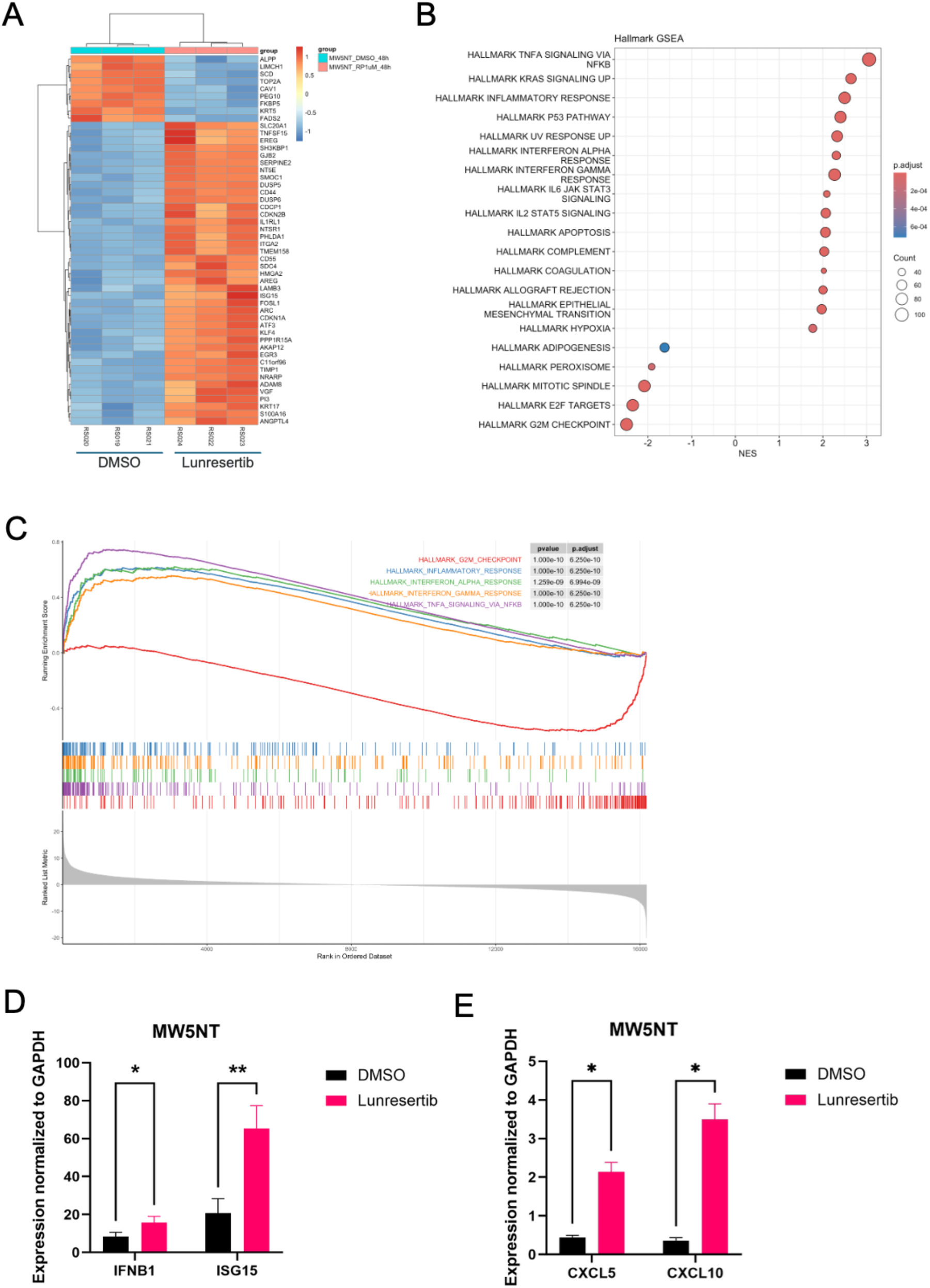
PKMYT1 inhibition induces pro-inflammatory signaling in *CCNE1-*amplified GEA. **A.** Heatmap of top differentially expressed genes in *CCNE1*-amplified MW5NT cells treated with vehicle or 1uM of lunresertib for 48H. **B.** HALLMARK pathway enrichment analysis on RNA-seq data of MW5NT treated with 1uM of lunresertib for 48H. **C.** Gene set enrichment analysis of the indicated pathways with plot showing the distribution of genes in the ranked list. **D.** *IFNB1* and *ISG15* levels in MW5NT treated with DMSO or 1μM of lunresertib for 48H quantified through qRT-PCR. **E.** *CXCL5* and *CXCL10* levels in MW5NT treated with DMSO or 1μM of lunresertib for 48H quantified through qRT-PCR.

### PKMYT1 inhibition demonstrates antitumor activity in *CCNE1*-overexpressing GC *in vivo*

To assess the antitumor activity of PKMYT1 inhibition *in vivo* and evaluate the functional implications of lunresertib-induced inflammatory signaling on the tumor-immune microenvironment, we employed a syngeneic xenograft model of *CCNE1*-amplified GC. Similar to the cytotoxicity observed in *CCNE1*-amplified human GEA cells, lunresertib exhibited *in vitro* activity against mouse GC organoids engineered to overexpress CCNE1 and lack p53 (CCNE1;*Trp53*^-/-^) (Figure 4A) (30). We validated that CCNE1;*Trp53*^-/-^ organoids had high CCNE1 expression by qRT-PCR (Supplementary Figure 3A). We injected murine CCNE1;*Trp53*^-/-^ GC organoids into the flanks of C57BL/6 mice to generate subcutaneous xenografts. Mice were randomized to receive either vehicle or lunresertib 15 mg/kg orally twice daily, which was well tolerated with no evidence of body weight loss (Supplementary Figure 3B). We observed that tumor growth was inhibited in mice administered lunresertib compared to vehicle control (Figure 4B). Representative hematoxylin and eosin (H&E) images of harvested tumors are shown in Figure 4C. Using immunofluorescence, we found that cleaved caspase 3 levels were higher in lunresertib-treated xenografts compared to vehicle controls, indicating that PKMYT1 inhibition promoted cell death via apoptosis (Figure 4D, E; Supplementary Figure 4). In addition, Ki67 expression, a marker of cellular proliferation, decreased in CCNE1;*Trp53*^-/-^ GC organoid xenografts in lunresertib-treated mice (Figure 4D, E; Supplementary Figure 4). Consistent with the effect of lunresertib treatment on human *CCNE1*-amplified GEA cells *in vitro*, lunresertib-treated tumors also demonstrated higher γH2AX expression, suggesting that lunresertib results in increased DNA damage in *CCNE1*-amplified GC *in vivo* (Figure 4D, E; Supplementary Figure 4). Overall, these data show that lunresertib has robust antitumor activity in *CCNE1*-amplified GC *in vivo*, consistent with other CCNE1-overexpressing preclinical models.

**Figure 4.**
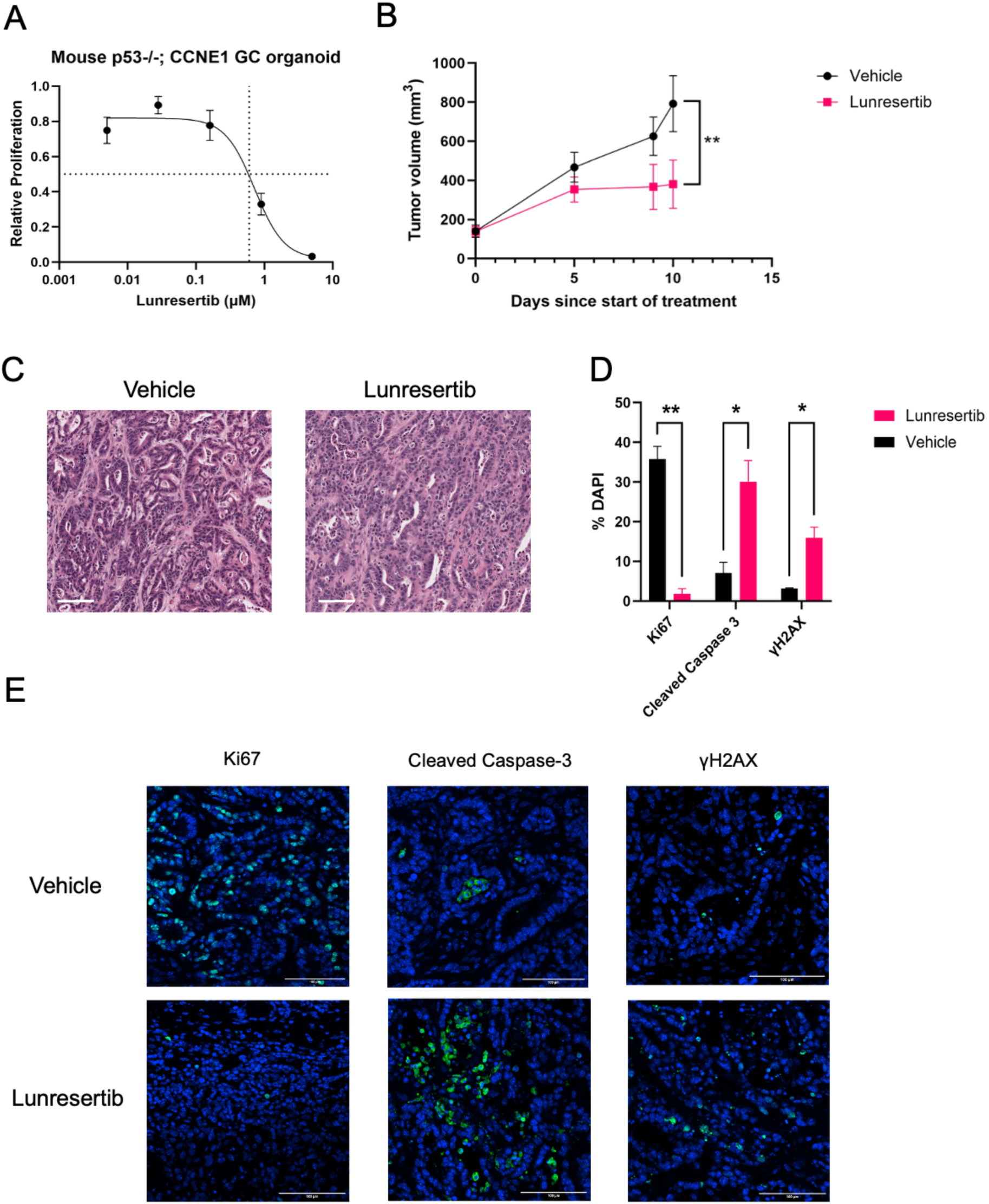
PKMYT1 inhibition demonstrates antitumor activity in *CCNE1*-overexpressing GC *in vivo*. **A.** Relative proliferation of CCNE1;*Trp53*^-/-^mouse GC organoids treated with indicated concentrations of lunresertib**. B.** Tumor volume (mean ± SEM) of CCNE1;*Trp53*^-/-^ xenograft models treated with vehicle or lunresertib (15mg/kg PO BID 5 days on/ 2days off). **C.** Representative H&E staining of CCNE1;*Trp53*^-/-^ tumor xenografts treated with vehicle or lunresertib. **D-E.** Ki67, Cleaved Caspase-3, and γH2AX expression in vehicle and lunresertib-treated tumors by immunofluorescence staining. *p < 0.05; **p <0.01, Welch’s t test.

### PKMYT1 inhibition modulates the tumor-immune microenvironment of *CCNE1*-overexpressing GC *in vivo*

Because lunresertib treatment elicited strong induction of inflammatory cytokines such as *IFNB1*, *ISG15*, and *CXCL10* in *CCNE1*-amplified GEA cells *in vitro*, we postulated that PKMYT1 inhibition may shape the tumor microenvironment (TME) and promote anti-tumor immunity. To test this, we stained tumor samples from mice treated with either vehicle or lunresertib using various immune cell markers and assessed immune cell infiltration by immunofluorescence. Lunresertib-treated tumors showed a marked increase in CD45, CD4, and CD8a positive cells, suggesting that PKMYT1 inhibition can promote accumulation of helper T cells as well as tumor-infiltrating cytotoxic T cells (Figure 5A-C; Supplementary Figure 5). In contrast, CD11b levels were decreased in lunresertib-treated mice, suggesting lower presence of myeloid cells which are often associated with immunosuppressive functions within the TME (Figure 5D; Supplementary Figure 5). These findings indicate that PKMYT1 inhibition can remodel the TME of *CCNE1-*amplified GC, promote T cell infiltration and activation, and shift the tumor milieu from an immune-cold to an immune-hot state.

**Figure 5.**
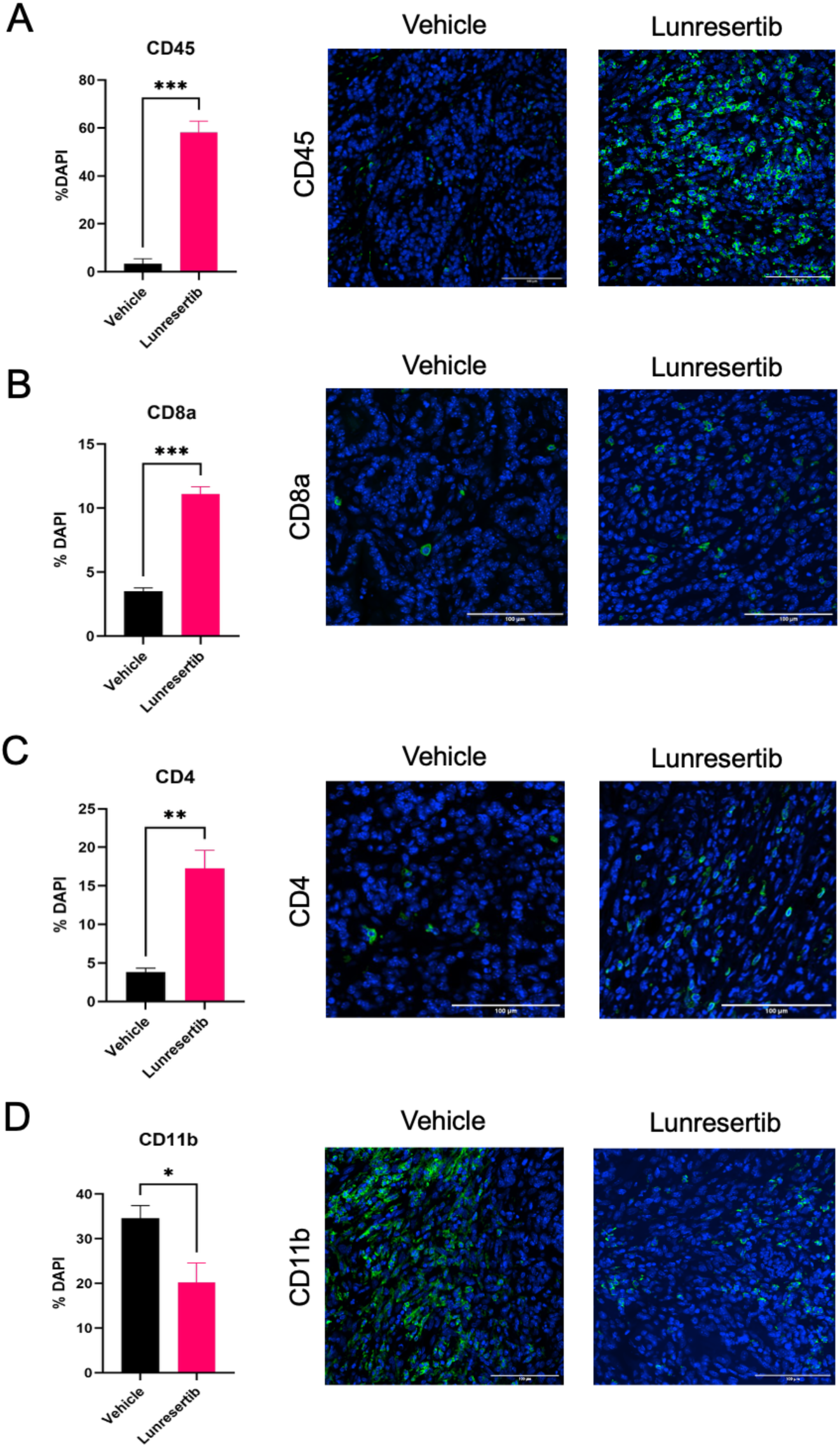
PKMYT1 inhibition modulates the tumor-immune microenvironment of *CCNE1*-overexpressing GC *in vivo*. **A-D.** Immunofluorescence staining of CD45, CD8a, CD4, and CD11b on lunresertib and vehicle treated tumor tissue with representative images. *p < 0.05, Welch’s t test.

### PKMYT1 inhibition synergizes with immune checkpoint inhibition

We hypothesized that the greater immune cell infiltration and pro-inflammatory changes in the TME induced by lunresertib could potentiate the efficacy of immune checkpoint inhibition (ICI) in *CCNE1*-amplified GC. To test this, syngeneic CCNE1;*Trp53*^-/-^ murine GC organoids were implanted subcutaneously into C57BL/6 mice to generate flank tumors and were randomized to receive vehicle, lunresertib, anti-PD-1, or the combination of lunresertib and anti-PD-1. Lunresertib and vehicle were administered orally 15 mg/kg twice daily, and anti-PD-1 was administered on day 1, 4, 6 of treatment. Both lunresertib and anti-PD-1 monotherapy moderately reduced tumor growth relative to vehicle control (Figure 6A, B). By contrast, the combination of lunresertib and anti-PD-1 showed the most pronounced anti-tumor activity, with deep tumor regressions compared to baseline tumor volumes (Figure 6A, B). Thus, in a preclinical model of *CCNE1*-amplified GC, PKMYT1 inhibition synergized with ICI to elicit robust anti-tumor responses.

**Figure 6.**
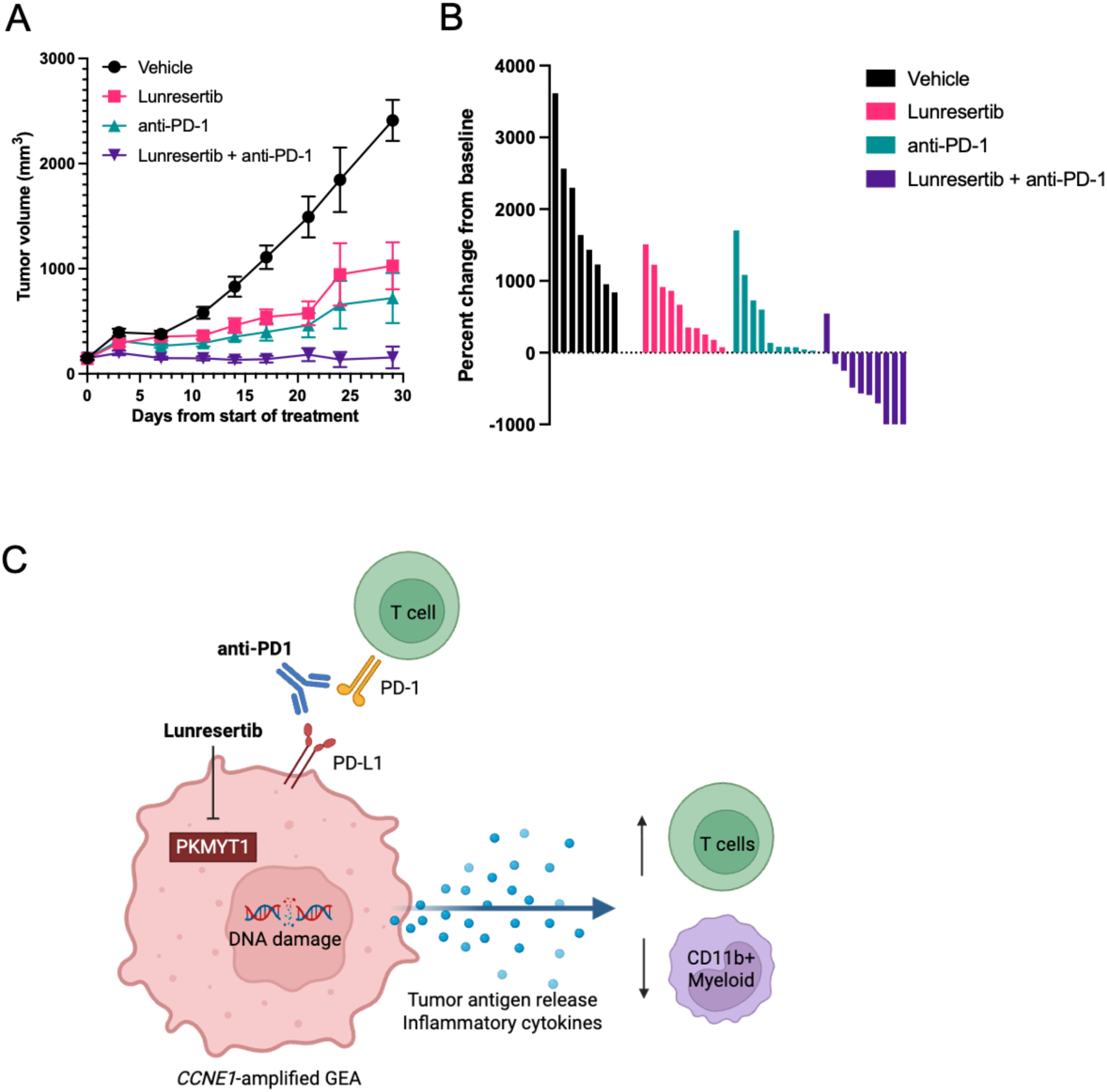
PKMYT1 inhibition synergizes with immune checkpoint inhibition. **A**. Tumor growth of CCNE1;*Trp53*^-/-^ xenograft models treated with vehicle, lunresertib, anti-PD-1, or lunresertib combined with anti-PD-1 for 30 days. Vehicle and lunresertib were administered daily (15mg/kg PO BID 5 days on/ 2 days off) and anti-PD-1 (200µg) on day 1, 4, 6**. B.** Waterfall plot showing percent change of tumor volume from baseline. **C.** Proposed mechanism of lunresertib and anti-PD-1 combination therapy.

## Discussion

*CCNE1* amplification is a key driver of CIN in GEA and is linked to therapeutic resistance, liver metastasis, and poor survival. Despite its clinical relevance, the role of *CCNE1* amplification as a potentially actionable in GEA has not been thoroughly explored. Here, using preclinical models of *CCNE1*-amplified GEA, we demonstrate that synthetic lethality with PKMYT1 inhibition represents a promising therapeutic strategy. Lunresertib treatment induces DNA damage and transcriptional upregulation of pro-inflammatory programs, including cytokines like *CXCL10* and type I interferon. *In vivo,* treatment of *CCNE1*-amplified tumor xenografts with lunresertib remodels the tumor microenvironment, enhancing cytotoxic T cell infiltration while reducing potentially immunosuppressive myeloid populations (Figure 6C). Moreover, combining PKMYT1 inhibition with PD-1 blockade produces synergistic antitumor effects, leading to tumor regression.

*CCNE1* amplification can promote CIN in part through persistent DNA replication stress. Mechanistically, replication errors during mitosis lead to excessive micronuclei, which frequently rupture and release cytosolic DNA (31). This leads to the chronic activation of the cGAS–STING (cyclic GMP-AMP synthase–stimulator of interferon genes) cytosolic DNA-sensing pathway and upregulation of PD-L1, enabling tumor cells to evade cytotoxic CD8+ T cells (32). Paradoxically, although cGAS-STING activation can promote an inflammatory state, the resulting TME remains immune-cold, characterized by poor CD8+ T cell infiltration, low levels of antigen expression, and the presence of immunosuppressive cell populations such as tumor-associated macrophages and T-regulatory cells (33). Such immune-cold TMEs are associated with resistance to targeted therapies, particularly ICIs. In accordance with this observation, translational analysis of tumor samples from the CheckMate-649 study, a randomized trial investigating first-line chemotherapy plus nivolumab compared to chemotherapy alone for metastatic GEA, found that CIN tumors derived less overall survival benefit with chemotherapy plus anti-PD-1 compared to other molecular subtypes (34). Similarly, *CCNE1*-amplified GEA has also been associated with decreased CD8+ T cell and B cell infiltration together with increased myeloid populations (10). Therefore, CIN that is driven by *CCNE1* amplification may suppress effective anti-tumor immunity, underscoring the need for therapeutic strategies that can reprogram this immunosuppressive TME.

One approach to enhance ICI efficacy in immune-cold tumors is to modulate the innate DNA Damage Response (DDR), thereby stimulating an immune response against evasive tumors (35). In *CCNE1*-amplified cells, PKMYT1 inhibition with lunresertib induces unscheduled CDK1 activation, resulting in premature entry into mitosis, replication stress, and eventually cell death. In this study, we validated that lunresertib selectively targets *CCNE1*-amplified GEA cells, induces DNA damage, and activates inflammatory signaling. This DDR-driven stress response remodels the TME by promoting cytotoxic T cell infiltration while decreasing immunosuppressive myeloid cell populations. Utilizing the *CCNE1*-amplified GC model as well as other preclinical models, other studies have also shown that depleting immunosuppressive myeloid cells such as myeloid-derived suppressor cells can also boost immunotherapy responses (36). We therefore tested the hypothesis that ICIs could amplify the immune activation initiated by lunresertib, and we found that combining PKMYT1 inhibition with anti-PD1 produced synergistic anti-tumor effects.

Other studies have explored similar strategies of targeting the DDR pathway in conjunction with immunotherapy. For example, one promising therapeutic target is Ataxia telangiectasia and Rad3-related (ATR) kinase, which regulates cell division and safeguards genomic integrity through arrest of cell cycle-progression and stabilization of stalled replication forks to enable DNA repair (37–39). ATR inhibition can lead to accumulation of DNA breaks, premature mitosis, and cell death, especially in tumors with defective DNA repair such as ATM loss. Preclinical studies have shown that ATR inhibitors can promote cytotoxic T cell accumulation and function, as well as enhance PD-1/PD-L1 blockade (40,41). This strategy of combined ATR inhibition and anti-PD-L1 therapy is being evaluated in clinical trials, such as the phase II HUDSON trial for immunotherapy-refractory advanced non-small-cell lung cancer (NSCLC) (NCT03334617), which has demonstrated encouraging efficacy with the combination of ceralasertib and durvalumab (42). Likewise, Wee1 inhibitors have also been shown to augment anti-tumor immune responses in preclinical models (43). Our findings suggest that combining lunresertib with immune checkpoint blockade may represent a rational, biomarker-driven strategy that leverages the selective vulnerability of *CCNE1*-ampliflication.

While lunresertib treatment induced inflammatory transcriptional programs in *CCNE1*-amplified GEA, the precise mechanisms by which lunresertib activates innate immune pathways and synergizes with immune checkpoint blockade remains to be further defined. Prior studies of DDR inhibition have shown that accumulation of cytosolic DNA can engage innate immune sensing pathways like cGAS-STING, resulting in type I interferon upregulation. Beyond cGAS-STING, additional pathways may also contribute. For example, our transcriptional profiling revealed significant enrichment of NFκB and JAK-STAT signaling following lunresertib exposure. Moreover, although we found increased infiltration of immune cells including cytotoxic T cells alongside decreased CD11b+ myeloid cells in lunresertib-treated tumors, the specific immune subsets responsible for mediating these anti-tumor effects remain to be determined.

Lunresertib is currently being evaluated in early phase clinical trials for solid tumors harboring *CCNE1* amplification, as well as genomic alterations that can lead to CCNE1 overexpression or sensitize to PKMYT1 inhibition such as pathogenic *FBXW7* or *PPP2R1A* mutations (NCT04855656, NCT05147272, NCT05147350). While lunresertib monotherapy was well-tolerated and led to some clinical responses, the efficacy appears to be overall modest, suggesting that combination therapy is warranted to enhance activity (44). Although lunresertib has not been investigated in combination with immunotherapy clinically, early-phase trials have explored combinations with the ATR inhibitor camonsertib, based on strong preclinical evidence suggesting synergy. ATR plays a crucial role in stabilizing DNA replication forks, which can be activated from replication stress driven by *CCNE1* amplification and PKMYT1 inhibition. The combination of PKMYT1 and ATR inhibition was shown to induce lethal levels of DNA damage and apoptosis, and preclinical data demonstrate strong efficacy across biomarker and tumor subtypes, especially in ovarian and endometrial cancers where these alterations are enriched (27). Therefore, an alternative strategy is pairing lunresertib with other agents targeting the DDR pathway in GC in combination with immunotherapy. In addition, while lunresertib is a first-in-class inhibitor, several other PKMYT1 inhibitors are also in preclinical or clinical development.

Our study does have some limitations. For example, our syngeneic mouse model may not fully recapitulate human GEA, including the heterogeneity, immune environment, and molecular features found in patient tumors. It is also unknown whether PKMYT1 inhibition may have additional effects on other cell types besides CCNE1 overexpressing tumor cells. The precise molecular and cellular mechanisms by which PKMYT1 inhibition reshapes the TME and potentiates anti-tumor immunity remains to be determined, which will be the focus of future studies.

In summary, our study shows that PKMYT1 inhibition in *CCNE1*-amplified GEA promotes antitumor activity by inducing DNA damage and modulating the tumor immune microenvironment. By eliciting a strong inflammatory transcriptional program, lunresertib may enhance cytotoxic T-cell infiltration and reduce immunosuppressive myeloid cells. This more inflamed TME may be more conducive to immunotherapy, resulting in stronger antitumor activity when combined with PD-1 blockade. Together, these insights provide strong rationale for clinical investigation of targeted combination therapies as a strategy to improve outcomes in *CCNE1*-amplified GEA. As many other tumors also harbor *CCNE1* amplification, this strategy of combined PKMYT1 inhibition and immunotherapy may also be relevant to other cancer types.

## Methods

### Cell culture

*CCNE1*^amp^ lines (copy number [CN] > 5) MW5NT and MW5FJ were obtained from Broad Institute of MIT and Harvard. SNU668 was obtained from the Korean Cell Line Bank. MW5NT and MW5FJ cells were cultured in Conditionally Reprogrammed Cells “F” Media (FM) made according to Cancer Cell Line Factory SOP: DMEM/F12 with NaPyr (Life Technologies 11330-032), DMEM with NaPyr (Life Technologies 11995-065), FBS (Life Technologies 26140-079), EGF Life Technologies PHG0311L), Insulin (Thermo 12585014-5mL), Hydrocortisone (Sigma H0888-1G), Cholera Toxin (Sigma C-8052-2MG), Y-27632 (Enzo ALX-270-333-M025), Pen/Strep/Glut (100x) (Life Technologies 10378-016), Fungizone (250 µg/mL) (Invitrogen 15290018), Gentamicin (50 mg/mL) (Life Technologies 15750-060). SNU668 cells were cultured in RPMI1640 (Gibco) with 10% fetal bovine Serum (FBS; Thermo Fisher) and 1% penicillin/streptomycin (P/S; Thermo Fisher). CCNE1;*Trp53*^-/-^ organoids were obtained from Adam Bass lab and cultured in Matrigel (Corning 354234) with conditioned media containing 50% L-WRN, 29% DMEMF/12 with NaPyr (Life Technologies 11330-032), 20% FBS (Life Technologies 26140-079), 1% Pen/Strep/Glut (100x) (Life Technologies 10378-016).

### In vitro cytotoxicity assays

Cells were plated onto 96-well plates at 10000 cells/well and incubated overnight. Cells were then treated with control (DMSO) or lunresertib at indicated conditions for 3 days and assessed for cell proliferation using the WST-1 assay. Briefly, 10 μL of WST-1 mixture (Cayman Chemical, Ann Arbor, MI) were added to each well and mixed for one minute on an orbital shaker. The cells were then incubated for 2 hours at 37 °C and absorbance was measured on a microplate reader (Molecular Devices SpectraMax iD3) at a wavelength of 450 nm.

### Clonogenic colony formation assays

Cells were plated onto 6-well plates at 5000 cells/well and cultured overnight in triplicate. They were then treated with control (DMSO) or lunresertib at indicated conditions every 3 days until the control group became confluent for a total of 14 days (SNU668) or 28 days (MW5NT). Cells were then fixed and stained with 0.1% Crystal violet in 20% methanol solution. The plates were washed, air-dried, scanned, and quantified in ImageJ (National Institutes of Health, Bethesda, MD).

### Western blotting

Cells were treated and collected at the indicated conditions, then washed and lysed with RIPA Buffer (Thermo Scientific) containing a protease and phosphatase (Millipore Sigma) inhibitor cocktail. Protein concentration was measured using the BCA Kit (Thermo Scientific) and incubated with NuPAGE™ LDS Sample Buffer (4X) (Invitrogen) containing 1:10 ratio of 2-Mercaptoethanol (Bio-Rad) at 95 °C for 5 minutes. Standardized protein samples were separated on 4-12% SDS-PAGE gels, electrotransferred to nitrocellulose membrane (Bio-Rad), blocked with 4% milk Bio Rad Blotting Grade Blocker Non-Fat Dry Milk (Bio-Rad) in Phosphate Buffered Saline-Tween (Boston BioProducts) (1x PBST), and immunoblotted with respective primary antibodies including: cyclin E (sc-247), mouse anti-Histone H2AX (MA1-2022). Membranes were then washed and blotted with species-appropriate horseradish peroxidase conjugated m-IgG Fc (sc-525409) in 4% milk in 1x PBST for 1 h, followed by chemiluminescent substrate (Thermo Scientific) incubation. Immunoblots were visualized using iBright™ CL1500 Imaging System (Invitrogen).

### Quantitative RT-PCR

Total RNA was extracted from cells at indicated conditions using Direct-zol RNA Miniprep Kits (Zymo Research) and quantified using Thermo Scientific NanoDrop Lite Plus Spectrophotometer. Reverse transcription was performed using TaqMan™ MicroRNA Reverse Transcription Kit (Applied Biosystems) according to manufacturer’s instructions. RT-PCR was performed in a total volume of 10 μL containing 1 μL of cDNA, 2 μL of primers, 5 μL of SYBR™ Green Universal Master Mix (Applied Biosystems), and 2 μL of nuclease free water (Fisher Scientific). The program steps for amplification were 95 °C for 2 minutes, 40 cycles of 95 °C for 15 s, and 60 °C for 1 minute. Relative mRNA expression was calculated using the 2-ΔΔCT method and GAPDH as an endogenous control. The primers were synthesized by Integrated DNA Technologies and the primer sequences are listed in Supplemental Table 1.

### Immunofluorescence

Cells were plated onto 8-well Nunc Lab-Tek II Chamber Slide System (Thermo Scientific) at 5000 cells/well and treated with control (DMSO) or lunresertib at indicated conditions and incubated for 48 hours. The cells were then fixed, permeabilized with 0.5% TritonX-100 (TX-100) in PBS, blocked with 1% BSA with 0.3% TX-100 in PBS at room temperature for 1 hour, and incubated overnight in 4 °C with primary antibodies: mouse anti-Histone H2AX (1:200, MA1-2022, Invitrogen). The slides were then washed and incubated with secondary antibodies: donkey anti-mouse IgG H&L (Alexa Fluor® 594), 1:500, Abcam. Slides were then imaged using Nikon Ti2 inverted microscope with AXR.

Flank tumors were harvested and fixed in formalin for 24H and transferred to 70% ethanol. Tumor tissues were then processed by Columbia University Irving Medical Center Molecular Pathology/MPSR (HICCC) core to generate formalin-fixed paraffin-embedded (FFPE) slides. Immunofluorescence staining was performed on the FFPE slides with antibodies listed in Supplemental Table 2 and imaged using Nikon Ti2 inverted microscope with AXR.

### Micronuclei staining

For micronuclei staining cells were plated on an opaque, clear-bottom 96-well plate at 10000 cells/well in triplicate and treated with control (DMSO) or lunresertib at indicated conditions and incubated for 48 hours. The cells were then fixed, permeabilized with 10% TX-100 in PBS, and incubated overnight in 4 °C with Hoechst 33342 (1:5000, 62249, Thermo Scientific) in PBS. The cells were then washed and imaged using ZEISS Celldiscoverer 7.

### RNA sequencing

Total RNA was extracted from cells at indicated conditions using Direct-zol RNA Miniprep Kits (Zymo Research) and quantified using Agilent Bioanalyzer 2100 through the Molecular Pathology Shared Resource (MPSR) at Columbia University (New York, NY). All library preparation and next-generation sequencing of total RNA was performed by JP Sulzberger Columbia Genome Center (New York, NY). STRPOLYA libraries were generated using a poly-A pulldown for mRNA, and pooled samples were sequenced on Illumina NextSeq 500/550. The R package DESeq2 (v1.46.0) was used to identify differentially expressed genes, and the gene-level counts were modeled using a negative binomial generalized linear model in DESeq2, with size-factor normalization and gene-wise dispersion estimation. Wald statistics and p-values were adjusted for multiple testing using the Benjamini–Hochberg false discovery rate. MsigDB (v24.1.0) was used to perform gene set enrichment analysis (GSEA). Additionally, pheatmap (v1.0.13), EnhancedVolcano (v1.24.0), enrichplot (1.26.6), and ggplot2 (v3.5.2) were used to generate the plots.

### Animal studies

All mice experiments were performed in adherence to the policies of the NIH Guide for the Care and Use of Laboratory Animals and approved by the Institutional Animal Care and Use Committee (IACUC). C57BL/6J mice (The Jackson Laboratory) aged 6 to 10 weeks were used for xenograft experiments. Mice were housed in an environmentally monitored, well-ventilated room maintained at a temperature of (23 ± 3 °C) and relative humidity of 40% - 70%, were exposed to 12 hours of light and dark cycles and were supplied with food and water ad libitum until reaching 6 weeks of age. Mice were injected subcutaneously with tumor cells in 100 μL of PBS and Matrigel mixture (1:1 ratio) in the right flank with 2 x 10^6^ cells. Mice health was monitored daily, and caliper measurements began when tumors were palpable. Tumors were measured three times a week, and tumor volume (TV) was determined using the formula A × B^2^ × 0.52 in which A and B are long and short diameters of a tumor. Treatment started when the tumor volume reached an average volume of ∼200mm^3^, and based on tumor volume and body weight mice were randomized into groups with *n*=6. Lunresertib was prepared according to the formulation protocol provided by Repare Therapeutics and administered 15 mg/kg BID daily. Anti-pd-1 (clone RMP1-14) was prepared in USP saline solution and 200 µg were administered on day 1, 4, 6 via intraperitoneal injection. Tumor samples were collected at indicated time points.

### Statistical analysis

The GraphPad Prism (10.5.0) and R (v4.4.2) were used to perform all statistical tests and analyses. P-value < 0.05 was considered statistically significant. The unpaired Student’s t-test was used to analyze the difference between two groups. Welch’s t-test was used to analyze the difference between two groups for nonparametric data. One-way or two-way ANOVA test was used to compare difference between multiple groups.

## Contributions

Conceptualization: AJB, RHM; Methodology: RHM; Investigation: SJ, LWW, SH, JP, CS, LLC, BI, AJB, RHM; Formal Analysis: SJ, LWW, SH, JP, CS, LLC, RHM; Supervision: RHM; Writing—original draft: SJ, LWW, JP, RHM; Writing—review and editing: SJ, LWW, SH, JP, CS, LLC, BI, AJB, RHM.

## Acknowledgements

LWW is supported by the NIH/NCI Molecular Oncology Training Program (5T32CA203703-09). RHM is supported by grants from the NIH (K08CA263304), Gastric Cancer Foundation, the 2023 AACR-Debbie’s Dream Foundation Innovation and Discovery Grant (Grant Number 23-80-41-MOY), and the US Department of Defense (HT9425-24-1-0419). This publication was supported by the NIH/NCI Cancer Center Support Grant P30CA013696. Lunresertib was provided by Repare Therapeutics.

---

**Supplemental Table 1.**
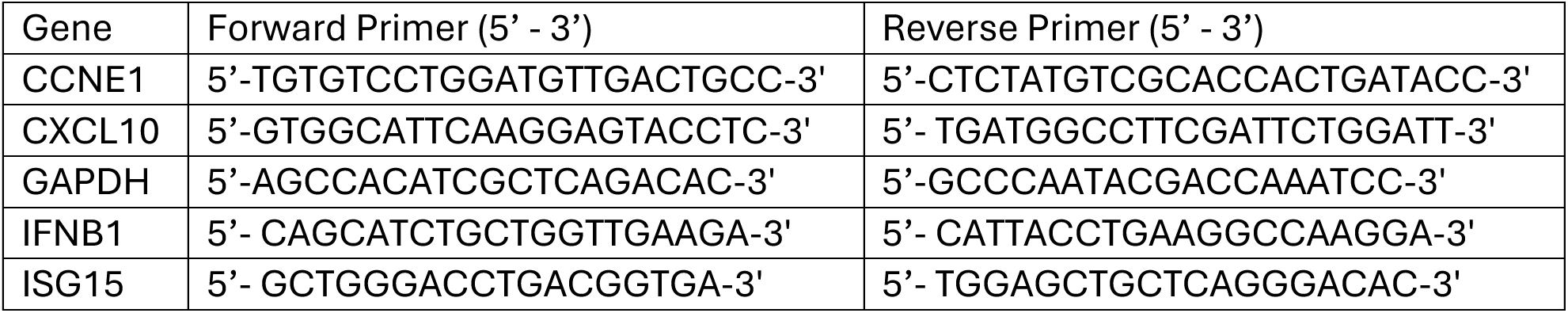

**Supplemental Table 2.**

| Antibody | Catalog |
| --- | --- |
| CD3 | Invitrogen 14-0032-82 |
| CD11b | Abcam ab133357 |
| CD19 | Biolegend 152402 |
| CD45 | Novus Bio NB100-77417SS |
| CD8a | Biolegend 100702 |
| Cleaved Caspase-3 | Cell Signaling 9661S |
| γH2AX | Abcam ab2893 |
| Ki67 | Abcam ab16667 |
| Ly6G | eBio 13-5931-82 |
| Goat anti-Rat IgG (H+L) Cross-Adsorbed Secondary Antibody, Alexa Fluor™ 488 | Invitrogen A-11006 |
| Donkey anti-Rabbit IgG (H+L) Highly Cross-Adsorbed Secondary Antibody, Alexa Fluor™ Plus 488 | Invitrogen A32790TR |

## References

1. Bray F, Laversanne M, Sung H, Ferlay J, Siegel RL, Soerjomataram I, et al. Global cancer statistics 2022: GLOBOCAN estimates of incidence and mortality worldwide for 36 cancers in 185 countries. CA Cancer J Clin. 2024;74(3):229–63.

2. Cancer Genome Atlas Research Network. Comprehensive molecular characterization of gastric adenocarcinoma. Nature. 2014 Sept 11;513(7517):202–9.

3. Cancer Genome Atlas Research Network, Analysis Working Group: Asan University, BC Cancer Agency, Brigham and Women’s Hospital, Broad Institute, Brown University, et al. Integrated genomic characterization of oesophageal carcinoma. Nature. 2017 Jan 12;541(7636):169–75.

4. Wu LW, Jang SJ, Shapiro C, Fazlollahi L, Wang TC, Ryeom SW, et al. Diffuse Gastric Cancer: A Comprehensive Review of Molecular Features and Emerging Therapeutics. Target Oncol. 2024 Nov;19(6):845–65.

5. Sundar R, Nakayama I, Markar SR, Shitara K, van Laarhoven HWM, Janjigian YY, et al. Gastric cancer. Lancet Lond Engl. 2025 June 7;405(10494):2087–102.

6. Shitara K, Ajani JA, Moehler M, Garrido M, Gallardo C, Shen L, et al. Nivolumab plus chemotherapy or ipilimumab in gastro-oesophageal cancer. Nature. 2022 Mar;603(7903):942–8.

7. Rha SY, Oh DY, Yañez P, Bai Y, Ryu MH, Lee J, et al. Pembrolizumab plus chemotherapy versus placebo plus chemotherapy for HER2-negative advanced gastric cancer (KEYNOTE-859): a multicentre, randomised, double-blind, phase 3 trial. Lancet Oncol. 2023 Nov;24(11):1181–95.

8. Qiu MZ, Oh DY, Kato K, Arkenau T, Tabernero J, Correa MC, et al. Tislelizumab plus chemotherapy versus placebo plus chemotherapy as first line treatment for advanced gastric or gastro-oesophageal junction adenocarcinoma: RATIONALE-305 randomised, double blind, phase 3 trial. BMJ. 2024 May 28;385:e078876.

9. Thrift AP, Wenker TN, El-Serag HB. Global burden of gastric cancer: epidemiological trends, risk factors, screening and prevention. Nat Rev Clin Oncol. 2023 May;20(5):338–49.

10. Rustgi N, Wu S, Samec T, Walker P, Xiu J, Lou E, et al. Molecular Landscape and Clinical Implication of CCNE1-amplified Esophagogastric Cancer. Cancer Res Commun. 2024 June 3;4(6):1399–409.

11. Cancer Genome Atlas Research Network. Comprehensive molecular characterization of gastric adenocarcinoma. Nature. 2014 Sept 11;513(7517):202–9.

12. Gorski JW, Ueland FR, Kolesar JM. CCNE1 Amplification as a Predictive Biomarker of Chemotherapy Resistance in Epithelial Ovarian Cancer. Diagn Basel Switz. 2020 May 5;10(5):279.

13. Zeng J, Hills SA, Ozono E, Diffley JFX. Cyclin E-induced replicative stress drives p53-dependent whole-genome duplication. Cell. 2023 Feb 2;186(3):528–542.e14.

14. Kok YP, Guerrero Llobet S, Schoonen PM, Everts M, Bhattacharya A, Fehrmann RSN, et al. Overexpression of Cyclin E1 or Cdc25A leads to replication stress, mitotic aberrancies, and increased sensitivity to replication checkpoint inhibitors. Oncogenesis. 2020 Oct 7;9(10):88.

15. Kim B, Shin HC, Heo YJ, Ha SY, Jang KT, Kim ST, et al. CCNE1 amplification is associated with liver metastasis in gastric carcinoma. Pathol Res Pract. 2019 Aug;215(8):152434.

16. Lee JY, Hong M, Kim ST, Park SH, Kang WK, Kim KM, et al. The impact of concomitant genomic alterations on treatment outcome for trastuzumab therapy in HER2-positive gastric cancer. Sci Rep. 2015 Mar 19;5:9289.

17. Bakhoum SF, Cantley LC. The Multifaceted Role of Chromosomal Instability in Cancer and Its Microenvironment. Cell. 2018 Sept 6;174(6):1347–60.

18. Davoli T, Uno H, Wooten EC, Elledge SJ. Tumor aneuploidy correlates with markers of immune evasion and with reduced response to immunotherapy. Science. 2017 Jan 20;355(6322):eaaf8399.

19. Thorsson V, Gibbs DL, Brown SD, Wolf D, Bortone DS, Ou Yang TH, et al. The Immune Landscape of Cancer. Immunity. 2018 Apr 17;48(4):812–830.e14.

20. Derks S, de Klerk LK, Xu X, Fleitas T, Liu KX, Liu Y, et al. Characterizing diversity in the tumor-immune microenvironment of distinct subclasses of gastroesophageal adenocarcinomas. Ann Oncol Off J Eur Soc Med Oncol. 2020 Aug;31(8):1011–20.

21. Gallo D, Young JTF, Fourtounis J, Martino G, Álvarez-Quilón A, Bernier C, et al. CCNE1 amplification is synthetic lethal with PKMYT1 kinase inhibition. Nature. 2022 Apr;604(7907):749–56.

22. Liu F, Stanton JJ, Wu Z, Piwnica-Worms H. The human Myt1 kinase preferentially phosphorylates Cdc2 on threonine 14 and localizes to the endoplasmic reticulum and Golgi complex. Mol Cell Biol. 1997 Feb;17(2):571–83.

23. Booher RN, Holman PS, Fattaey A. Human Myt1 is a cell cycle-regulated kinase that inhibits Cdc2 but not Cdk2 activity. J Biol Chem. 1997 Aug 29;272(35):22300–6.

24. Strausfeld U, Labbé JC, Fesquet D, Cavadore JC, Picard A, Sadhu K, et al. Dephosphorylation and activation of a p34cdc2/cyclin B complex in vitro by human CDC25 protein. Nature. 1991 May 16;351(6323):242–5.

25. Lang F, Kaur K, Zaheer J, Ribeiro DL, Yang C. Myt1 Kinase: An Emerging Cell-Cycle Regulator for Cancer Therapeutics. Clin Cancer Res Off J Am Assoc Cancer Res. 2025 Mar 17;31(6):960–4.

26. Takei S, Kawazoe A, Shitara K. The New Era of Immunotherapy in Gastric Cancer. Cancers. 2022 Feb 18;14(4):1054.

27. Xu H, George E, Gallo D, Medvedev S, Wang X, Datta A, et al. Targeting CCNE1 amplified ovarian and endometrial cancers by combined inhibition of PKMYT1 and ATR. Nat Commun. 2025 Apr 1;16(1):3112.

28. Li M, Lulla AR, Wang Y, Tsavaschidis S, Wang F, Karakas C, et al. Low-Molecular Weight Cyclin E Confers a Vulnerability to PKMYT1 Inhibition in Triple-Negative Breast Cancer. Cancer Res. 2024 Nov 15;84(22):3864–80.

29. Agustinus AS, Al-Rawi D, Dameracharla B, Raviram R, Jones BSCL, Stransky S, et al. Epigenetic dysregulation from chromosomal transit in micronuclei. Nature. 2023 July;619(7968):176–83.

30. Wu Z, Nagaraja A, Sun S, Lan L, Zhou J, Bass A. Abstract 2834: CyclinE1 amplification promotes chromosomal instability and distinct vulnerabilities in gastroesophageal adenocarcinoma. Cancer Res. 2025 Apr 21;85(8_Supplement_1):2834–2834.

31. Wilhelm T, Said M, Naim V. DNA Replication Stress and Chromosomal Instability: Dangerous Liaisons. Genes. 2020 June 10;11(6):642.

32. Yan Y, Tan X, Song B, Yi M, Chu Q, Wu K. Breaking barriers: The cGAS-STING pathway as a novel frontier in cancer immunotherapy. Cancer Commun Lond Engl. 2025 Nov;45(11):1513–46.

33. Wu B, Zhang B, Li B, Wu H, Jiang M. Cold and hot tumors: from molecular mechanisms to targeted therapy. Signal Transduct Target Ther. 2024 Oct 18;9(1):274.

34. Shitara K, Janjigian YY, Ajani J, Moehler M, Yao J, Wang X, et al. Nivolumab plus chemotherapy or ipilimumab in gastroesophageal cancer: exploratory biomarker analyses of a randomized phase 3 trial. Nat Med. 2025 May;31(5):1519–30.

35. Sen T, Rodriguez BL, Chen L, Corte CMD, Morikawa N, Fujimoto J, et al. Targeting DNA Damage Response Promotes Antitumor Immunity through STING-Mediated T-cell Activation in Small Cell Lung Cancer. Cancer Discov. 2019 May;9(5):646–61.

36. Qian J, Ma C, Waterbury QT, Zhi X, Moon CS, Tu R, et al. A CXCR4 partial agonist improves immunotherapy by targeting immunosuppressive neutrophils and cancer-driven granulopoiesis. Cancer Cell. 2025 Aug 11;43(8):1512–1529.e11.

37. Carrassa L, Damia G. DNA damage response inhibitors: Mechanisms and potential applications in cancer therapy. Cancer Treat Rev. 2017 Nov;60:139–51.

38. da Costa AABA, Chowdhury D, Shapiro GI, D’Andrea AD, Konstantinopoulos PA. Targeting replication stress in cancer therapy. Nat Rev Drug Discov. 2023 Jan;22(1):38–58.

39. Ngoi NYL, Pilié PG, McGrail DJ, Zimmermann M, Schlacher K, Yap TA. Targeting ATR in patients with cancer. Nat Rev Clin Oncol. 2024 Apr;21(4):278–93.

40. Taniguchi H, Chakraborty S, Takahashi N, Banerjee A, Caeser R, Zhan YA, et al. ATR inhibition activates cancer cell cGAS/STING-interferon signaling and promotes antitumor immunity in small-cell lung cancer. Sci Adv. 2024 Sept 27;10(39):eado4618.

41. Hardaker EL, Sanseviero E, Karmokar A, Taylor D, Milo M, Michaloglou C, et al. The ATR inhibitor ceralasertib potentiates cancer checkpoint immunotherapy by regulating the tumor microenvironment. Nat Commun. 2024 Feb 24;15(1):1700.

42. Besse B, Pons-Tostivint E, Park K, Hartl S, Forde PM, Hochmair MJ, et al. Biomarker-directed targeted therapy plus durvalumab in advanced non-small-cell lung cancer: a phase 2 umbrella trial. Nat Med. 2024 Mar;30(3):716–29.

43. Taniguchi H, Caeser R, Chavan SS, Zhan YA, Chow A, Manoj P, et al. WEE1 inhibition enhances the antitumor immune response to PD-L1 blockade by the concomitant activation of STING and STAT1 pathways in SCLC. Cell Rep. 2022 May 17;39(7):110814.

44. Yap TA, Schram A, Lee EK, Simpkins F, Weiss MC, LoRusso P, et al. Abstract PR008: MYTHIC: First-in-human (FIH) biomarker-driven phase I trial of PKMYT1 inhibitor lunresertib (lunre) alone and with ATR inhibitor camonsertib (cam) in solid tumors with CCNE1 amplification or deleterious alterations in FBXW7 or PPP2R1A. Mol Cancer Ther. 2023 Dec 1;22(12_Supplement):PR008–PR008.

