## Supplementary Figures for "PKMYT1 inhibition induces DNA damage and synergizes with immune checkpoint blockade in *CCNE1*-amplified gastroesophageal adenocarcinoma"

### Supplementary Figure 1

A

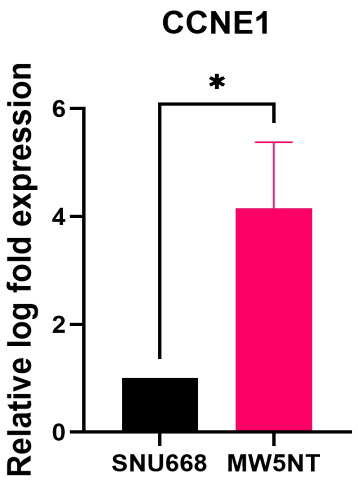


B

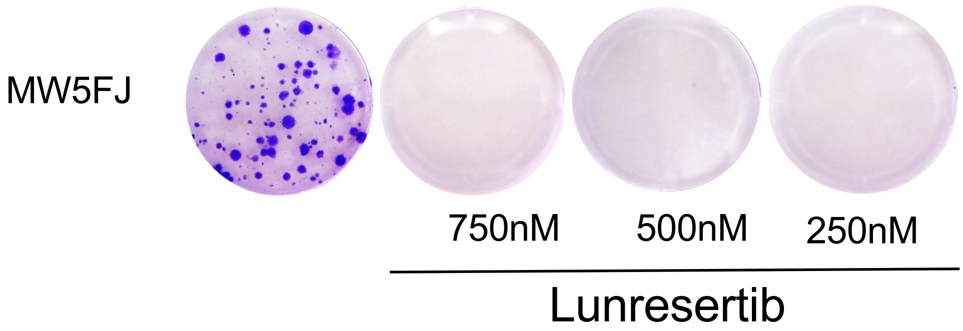


**Supplementary Figure 1. PKMYT1 inhibition selectively targets *CCNE1*-amplified GEA.**
**A.** *CCNE1*expression in *CCNE1*-amplified (MW5NT) and CCNE1 WT (SNU668) cell lines quantified by qRT-PCR. **B.**Clonogenic assay of *CCNE1*-amplified MW5FJ cells treated with indicated concentrations of lunresertib. *$p < 0.05$; unpaired t test.

### Supplementary Figure 2

A

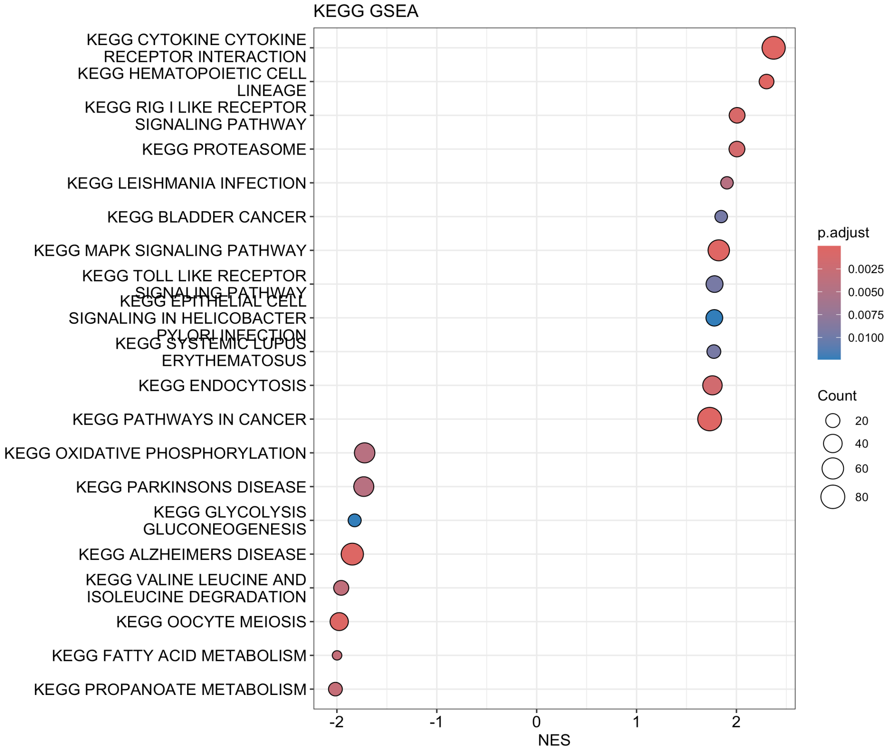


B

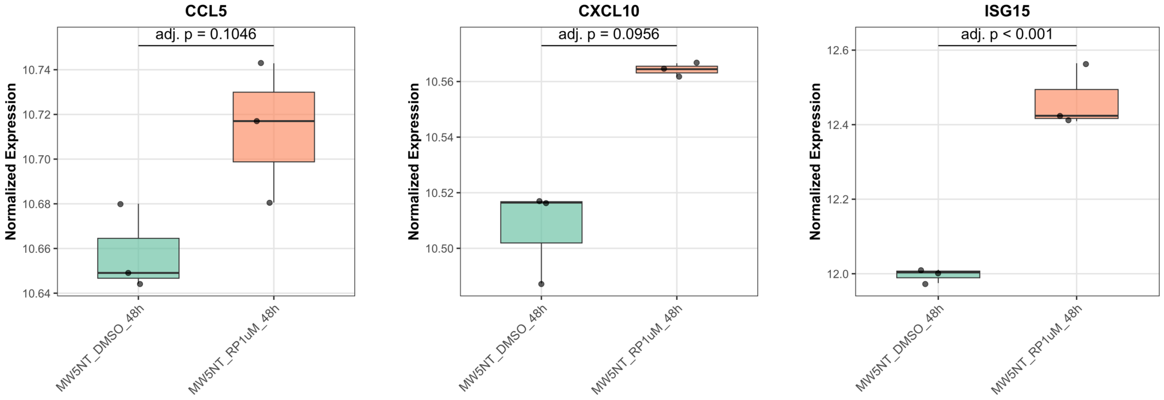


**Supplementary Figure 2. PKMYT1 inhibition induces pro-inflammatory signaling in *CCNE1*-amplified GEA.**
**A.** HALLMARK pathway enrichment analysis on RNA-seq data of MW5NT treated with 1uM of lunresertib for 48H. **B.** Box plot of *CCL5*, *CXCL10*, *ISG15* expression in MW5NT treated with vehicle or 1uM of lunresertib for 48H.

Supplementary Figure 3

A

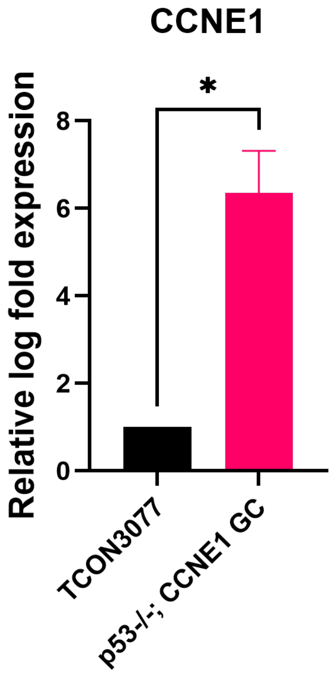


B

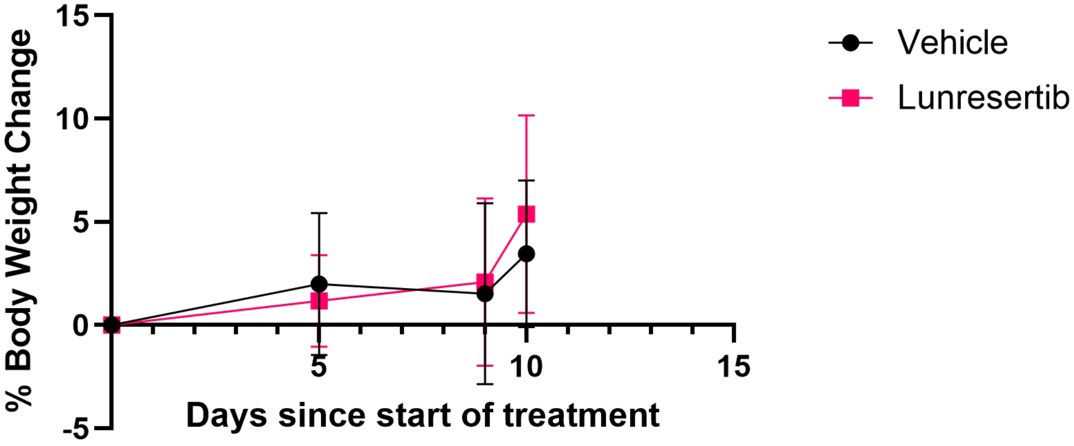


**Supplementary Figure 3. PKMYT1 inhibition demonstrates antitumor activity in *CCNE1*-overexpressing GC *in vivo.***
**A.** *CCNE1* expression in *CCNE1*-amplified (CCNE1;*Trp53*$^{-/-}$ GC) and CCNE1 WT (TCON3077) murine GC organoids quantified by qRT-PCR. **B.** Percent body weight change of mice since starting treatment of vehicle or lunresertib. *p < 0.05; Student's t test.

Supplementary Figure 4

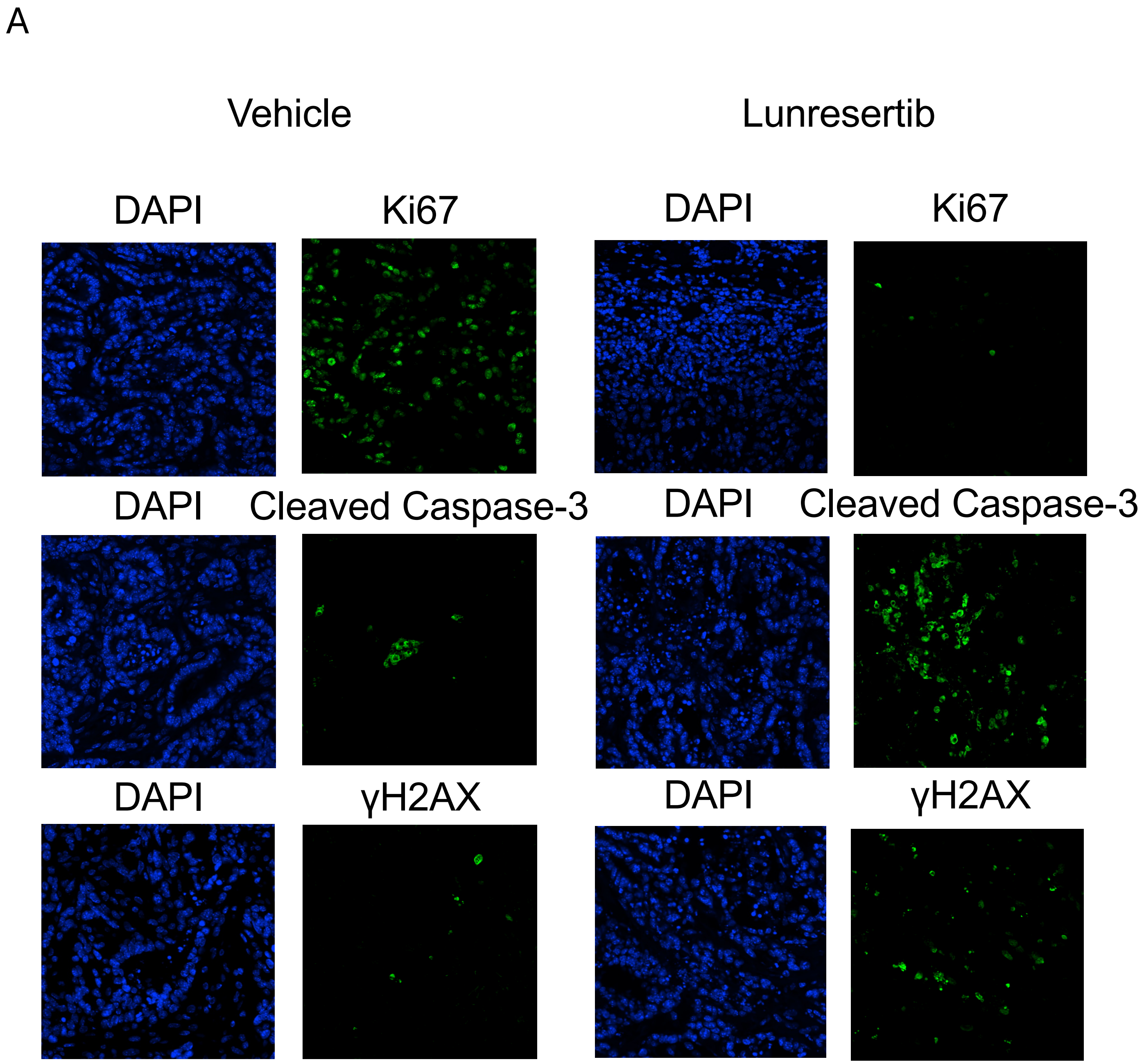


**Supplementary Figure 4. PKMYT1 inhibition demonstrates antitumor activity in *CCNE1*-overexpressing GC *in vivo.***

**A.** Immunofluorescence staining of Ki67, Cleaved Caspase 3, and γH2AX on lunresertib and vehicle treated tumor tissue with image panels of DAPI and respective antibodies.

### Supplementary Figure 5

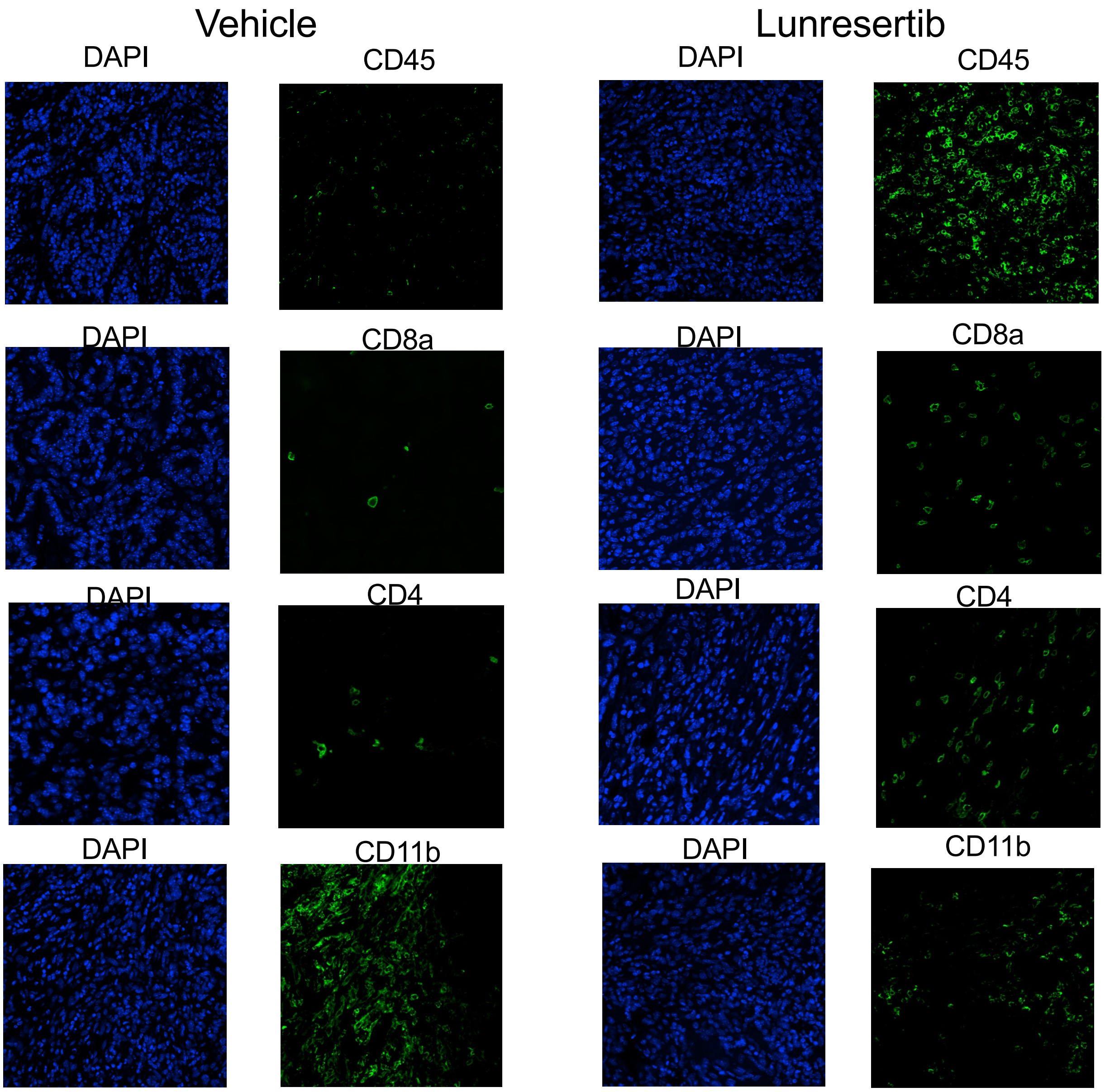


**Supplementary Figure 5. PKMYT1 inhibition modulates the tumor-immune microenvironment of *CCNE1*-overexpressing GC *in vivo.***

**A.** Immunofluorescence staining of CD45, CD8a, CD4, and CD11b on lunresertib and vehicle treated tumor tissue with image panels of DAPI and respective antibodies.
